# Exposure to auditory feedback delay while speaking induces perceptual habituation but does not mitigate the disruptive effect of delay on speech auditory-motor learning

**DOI:** 10.1101/2020.02.15.951004

**Authors:** Douglas M. Shiller, Takashi Mitsuya, Ludo Max

## Abstract

Perceiving the sensory consequences of our actions with a delay alters the interpretation of these afferent signals and impacts motor learning. For reaching movements, delayed visual feedback of hand position reduces the rate and extent of visuomotor adaptation, but substantial adaptation still occurs. Moreover, the detrimental effect of visual feedback delay on reach motor learning—selectively affecting its implicit component—can be mitigated by prior habituation to the delay. Auditory-motor learning for speech has been reported to be more sensitive to feedback delay, and it remains unknown whether habituation to auditory delay reduces its negative impact on learning. We investigated whether 30 minutes of exposure to auditory delay during speaking (a) affects the subjective perception of delay, and (b) mitigates its disruptive effect on speech auditory-motor learning. During a speech adaptation task with real-time perturbation of vowel spectral properties, participants heard this frequency-shifted feedback with no delay, 75 ms delay, or 115 ms delay. In the delay groups, 50% of participants had been exposed to the delay throughout a preceding 30-minute block of speaking whereas the remaining participants completed this block without delay. Although habituation minimized awareness of the delay, no improvement in adaptation to the spectral perturbation was observed. Thus, short-term habituation to auditory feedback delays is not effective in reducing the negative impact of delay on speech auditory-motor adaptation. Combined with previous findings, the strong negative effect of delay and the absence of an influence of delay awareness suggest the involvement of predominantly implicit learning mechanisms in speech.

**HIGHLIGHTS:**

- Speech auditory-motor adaptation to a spectral perturbation was reduced by ~50% when feedback was delayed by 75 or 115 ms.
- Thirty minutes of prior delay exposure without perturbation effectively reduced participants’ awareness of the delay.
- However, habituation was ineffective in remediating the detrimental effect of delay on speech auditory-motor adaptation.
- The dissociation of delay awareness and adaptation suggests that speech auditory-motor learning is mostly implicit.

## Introduction

When one or more sensory modalities (e.g., vision) are subjected to environmentally induced delays, motor control for tasks such as hand-writing can be severely disrupted (Morikiyo and Matsushima 1990; Smith et al. 1960; Tamada 1995). In fact, it has been shown that perceiving the sensory consequences of one’s actions with a delay as short as 100 ms can alter the interpretation of these afferent signals, causing them to be attributed to an external source rather than a self-generated action (Blakemore et al. 2000). It is therefore not surprising that the effects of feedback delay have also been explored in the context of sensorimotor *learning*: credit assignment for experienced errors is critical in these tasks given that sensorimotor learning requires the correct updating of internal representations of the association between motor commands and their sensory consequences in a particular environment (Shadmehr et al. 2010).

A number of previous sensorimotor learning studies have experimentally manipulated the visual feedback delay during reaching tasks in which participants wore prism glasses (Kitazawa et al. 1995; Tanaka et al. 2011) or in which the location of a computer cursor representing hand position was rotated around the center of the workspace (Honda et al. 2012a). Visual feedback representing only the final hand position at movement completion (Kitazawa et al. 1995; Tanaka et al. 2011) or the entire motion of the cursor (Honda et al. 2012a) was shown either with no delay (other than that inherent in the instrumentation setup, which by itself was sometimes as long as 60 ms) or after experimentally induced additional delays ranging from 50 to 10000 ms. In the absence of feedback delay, adaptation to these visual perturbations (i.e., prism displacement and visuomotor rotation) manifests as learned compensatory adjustments in movement planning such that participants aim in the opposite direction of the perturbation. Although visual feedback delays were found to negatively impact the initial rate of this adaptation (i.e., the slope of a learning function), the aforementioned studies all reported that a robust degree of visuomotor adaptation did occur by the end of training. With a delay of 100 ms, the final extent of prism adaptation was essentially unaffected in Tanaka et al. (2011) and reduced by approximately 35-40% in Kitazawa et al. (1995). With a delay of 200 ms, visuomotor rotation learning was reduced by approximately 20% in Honda et al. (2012a). Even with a much longer delay of 5000 ms, prism adaptation was not eliminated but reduced by approximately 50% in Kitazawa et al. (1995). Thus, while delaying visual feedback of hand position often had a statistically significant impact on the rate and extent of visuomotor adaptation^1^, the disruptive effect on the extent of learning has been limited, and substantial motor learning still occurs under those circumstances.

Interestingly, the potentially disruptive effects of delays in motor-related sensory signals may be reduced in part by neural mechanisms that can predict and compensate for such effects (Miall et al. 1993; Miall and Jackson 2006). Numerous studies have shown that sensory predictions related to self-generated movement take account of the temporal relationship between the motor and sensory processes and are modified through experience. Following prolonged exposure to a consistent time delay between an action such as a button press and a sensory consequence such as a tone, subjects begin to perceive the sensory event as shifted in time toward the action (Haggard et al. 2002; Haggard and Clark 2003; Park et al. 2003; Stetson et al. 2006; Heron et al. 2009). Similar perceived changes in the relative timing of sensorimotor signals have been observed in tasks involving visual or tactile feedback (Park et al. 2003; Stetson et al. 2006; Heron et al. 2009). Moreover, it has been demonstrated that the typical attenuation of neural and perceptual responses to self-generated sensory stimuli (as compared with external stimulation) also becomes delayed when participants consistently experience a delay between their movement and the sensory consequences of that movement (Aliu et al., 2009; Kilteni et al. 2019). Lastly, in a pair of studies examining perceptual habituation to auditory feedback delays during vocalization of individual vowels and connected speech, Yamamoto and Kawabata (2011, 2014) found that judgments of the simultaneity between motor act and auditory feedback shifted toward the physical delay (i.e., with exposure, delayed feedback became more likely to be judged as simultaneous with the production).

Most important for the present work, a small number of studies have examined whether such plasticity in the perception of the temporal relationship between motor actions and their sensory consequences can be leveraged for motor learning. In other words, can experience-based plasticity in the temporal aspect of sensory predictions negate the otherwise detrimental effects of feedback delays when learning to adapt to a different sensorimotor perturbation? Initially, this appeared to be not the case as Tanaka et al. (2011) found no improvement in prism glass adaptation with 136 ms visual feedback delay (100 ms added delay, 36 ms equipment delay) if subjects had also experienced the same delay before vision was shifted. However, in this Tanaka et al. (2011) study of pointing movements, (a) prior delay exposure was limited as it occurred only during the 60 baseline trials before the visual shift was implemented, (b) the subjectively perceived delay after this short period of exposure was still 96 ms, (c) the visual perturbation was introduced abruptly (thus, leading to subject awareness and explicit learning involving the use of cognitive strategies), and (d) the task was completed with only movement endpoint feedback rather than full trajectory feedback. In contrast, in studies of reaching movements completed with a 20-degree visual feedback rotation that was introduced gradually and with full motion path feedback, Honda (2012a, 2012b) tested whether the negative impact of a 260 ms visual feedback delay (200 ms added delay, 60 ms equipment delay) could be reduced by first habituating subjects to the delay for 100 or 120 movements, depending on the study. Findings revealed that subjects who experienced this amount of prior exposure to the delayed feedback showed improved learning in the subsequent adaptation task as compared with subjects without prior exposure to feedback delay. Thus, studies of sensorimotor adaptation in visually-guided arm movements indicate not only that delayed sensory feedback during a motor task results in consistent but limited effects on motor learning, but also that prior habituation to the feedback delay can mitigate these disruptive effects on visuomotor learning.

In contrast with visuomotor learning in the upper limb, the sensorimotor control of *speech articulation* appears to be more sensitive to feedback delays. Max and Maffett (2015) examined adaptation in subjects’ vowel production when the feedback signal was manipulated such that the frequencies of all resonance peaks (i.e., formants) were shifted upward, and this feedback signal was also delayed by 0, 100, 250, or 500 ms (in addition to a 10 ms delay inherent in the equipment). In the 0 ms delay condition, a robust adaptation effect was observed over 120 trials with formant-shifted auditory feedback: participants gradually learned to adjust the planning of their articulatory movements so that their formant frequencies changed in the opposite direction of the feedback shift, thereby partially cancelling the perturbation. However, with all delays of 100 ms or more, adaptation was completely absent. In a later study, Mitsuya et al. (2017) observed an 80-90% reduction in the extent of speech adaptation to altered feedback involving a shift of only the first formant when this signal was also delayed by 100 ms (although those authors reported that the remaining amount of adaptation was still statistically different from zero). Hence, as compared with visual feedback for upper limb movements, the processing of auditory feedback for the adaptive learning of oral speech movements may depend more strongly on a very tight temporal coupling between motor events and their sensory consequences.

It is plausible that this stronger effect of delayed feedback on sensorimotor learning for speech may relate to fundamental properties of the involved neural and biomechanical systems or more general characteristics of the underlying learning mechanisms. For example, evidence to date indicates that speech auditory-motor adaptation represents an almost entirely implicit form of learning. First, unlike standard visuomotor tasks, naive subjects are completely unaware of which vocal tract movement strategies can compensate for the implemented formant-shift perturbation (e.g., for a simultaneous upward perturbation of the first two formants F1 and F2, oral opening should be reduced to compensate in F1 and tongue protrusion should be reduced to compensate in F2 or smaller compensatory changes in both formants could be achieved through lip protrusion or rounding). Second, even when speech auditory-motor adaptation occurs in response to an abruptly introduced perturbation, subjects’ trial-by-trial reports indicate that they are unaware of having made any changes in their speech output (Kim and Max 2020). Third, there is no difference in the amount of speech adaptation to pitch-shifted auditory feedback in conditions where participants are instructed to either compensate or ignore the feedback (Keough et al. 2013) or in speech adaptation to formant-shifted auditory feedback conditions where participants are instructed to compensate, to ignore the feedback, or to explicitly avoid compensating (Munhall et al. 2003). Consequently, task differences in the involvement of implicit learning versus explicit strategy use may play a role in the differential effects of sensory delays on speech and limb adaptation. This suggestion is also consistent with recent work showing that, in upper limb motor learning, feedback delays negatively affect implicit learning but not explicit strategy selection (Brudner et al. 2016; McDougle and Taylor 2019; Schween and Hegele 2017). We therefore suggest that feedback delays may have only a relatively minor impact on visuomotor reach adaptation because it is characterized by a small implicit component and a large explicit component, at least in the case of an abruptly introduced perturbation (Anguera et al. 2010; Fernandez-Ruiz et al. 2011; McDougle et al. 2016; Taylor et al. 2014) or a gradually introduced perturbation with reward feedback (Holland et al. 2018). In contrast, the detrimental impact of feedback delays on auditory-motor speech learning may result from this type of learning being largely or exclusively implicit in nature as suggested by both pitch-shift and formant-shift experiments (Keough et al. 2013; Kim and Max 2020; Munhall et al. 2003).

If it is true that speech auditory-motor adaptation is entirely dependent on implicit learning mechanisms, then even prior habituation to an auditory feedback delay (eliminating participants’ explicit awareness of the delay) may not reduce the negative impact of delay on speech adaptation. As mentioned above, it has already been demonstrated that feedback delays selectively impair implicit learning (Brudner et al. 2016; Schween and Hegele 2017). We propose that in mixed implicit/explicit adaptation tasks where delay habituation largely restores the otherwise degraded learning (such as the visuomotor task in Honda 2012a, 2012b), habituated participants may achieve restored learning by increasing explicit strategy use when they subjectively no longer perceive the delay but nevertheless experience compromised task performance (due to the physical delay affecting implicit learning). If, on the other hand, a learning task were predominantly implicit in nature, even delay-habituated participants would not be able to adjust explicit strategy use to reduce performance error. We therefore investigated the presence versus absence of a facilitatory effect of delay habituation on speech auditory-motor learning by asking whether pre-exposure to a delay in the auditory feedback channel will (a) alter participants’ subjective perception of the imposed delay prior to and during completion of a formant-shift adaptation task, and (b) mitigate the previously documented disruptive effects of such a delay on the extent of speech auditory-motor adaptation.

## Experimental Procedures

### Overall study design

Fifty-five adults with no reported history of speech, hearing, or neurological disorders participated after providing written informed consent (all procedures were approved by the Institutional Review Board at the University of Washington). Each participant was pseudo-randomly assigned to a control group or one of four experimental groups so that each group included 5 men and 6 women (age information per group provided below). Based on a pure tone hearing screening procedure, fifty-four participants had thresholds at 20 dB HL or better for the octave frequencies from 250 Hz to 4 kHz (tested in both ears separately). The remaining participant had a threshold of 25 dB HL at 250 Hz for the left ear, but 20 dB HL in both ears for all other frequencies.

The study involved an initial *Delay Exposure* task (blocks of word reading and picture naming completed with or without feedback delay depending on group assignment) followed by an *Auditory-Motor Adaptation* task (blocks of word reading with formant-shifted auditory feedback completed with or without feedback delay depending on group assignment). Thus, the five groups differed in the amount of delay to which they were exposed in the adaptation task and whether or not they first habituated to this delay in the exposure task (Figure 1A):

1. no delay in the exposure task, no delay in the adaptation task (*Control Group*; age *M* = 24.09 years, *SD* = 5.70, range = 18-35)
2. no delay in the exposure task, 75 ms delay in the adaptation task (*No-Habituation Short-Delay Group*; age *M* = 22.64 years, *SD* = 5.90, range = 18-39)
3. no delay in the exposure task, 115 ms delay in the adaptation task (*No-Habituation Long-Delay Group*; age *M* = 23.27 years, *SD* = 6.10, range = 18-34)
4. 75 ms delay in the exposure task, 75 ms delay in the adaptation task (*Habituation Short-Delay Group*; age *M* = 24.82 years, *SD* = 6.40, range = 18-42)
5. 115 ms delay in the exposure task, 115 ms delay in the adaptation task (*Habituation Long-Delay Group*; age *M* = 24.18 years, *SD* = 4.47, range = 19-32)

**Figure 1.**
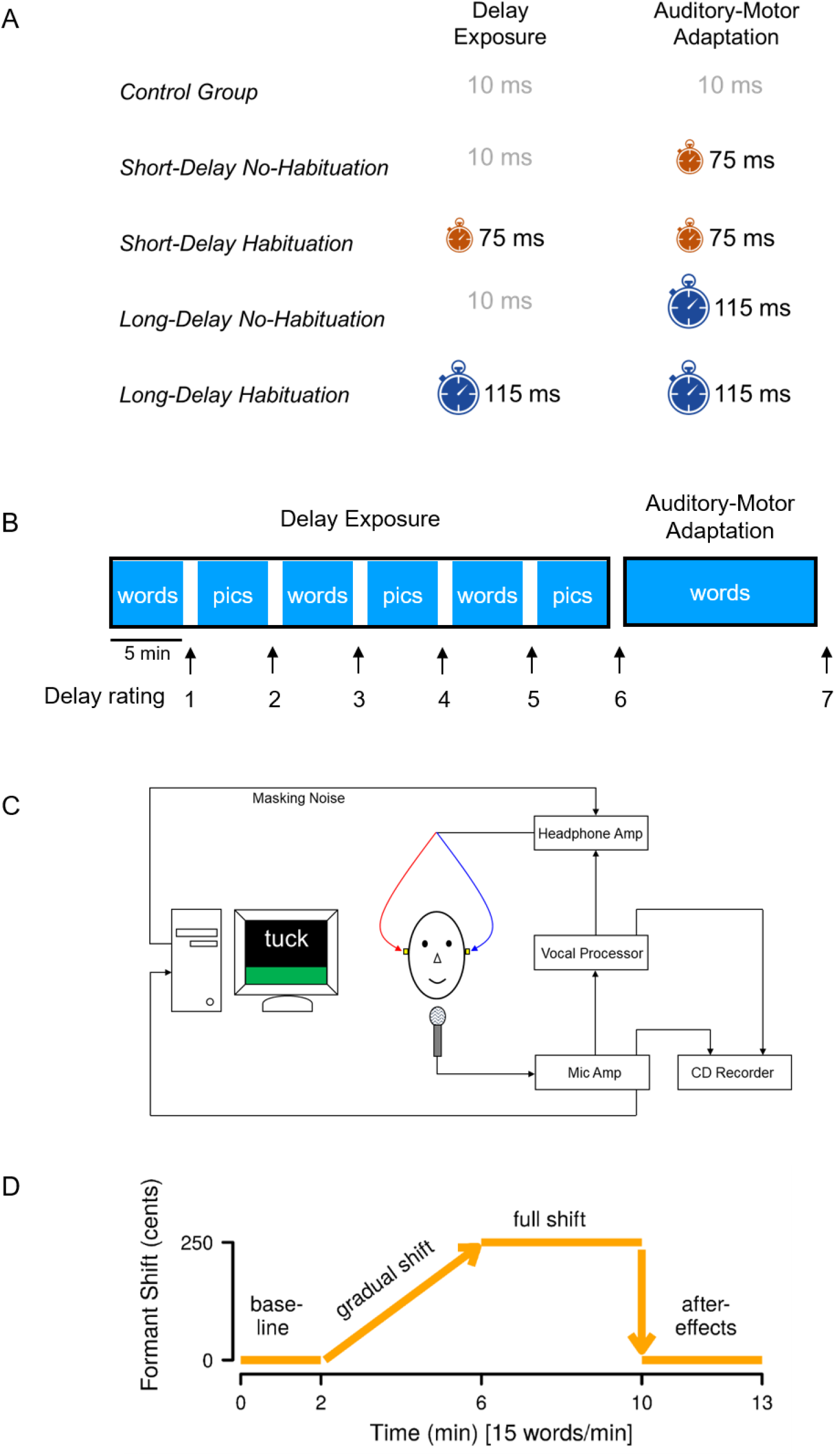
(A) Auditory feedback delay experienced by each of five subject groups during the Delay Exposure task and the subsequent Auditory-Motor Adaptation task. (B) Sequence of speech tasks completed in the Delay Exposure task (alternating blocks of word reading and picture naming while exposed to delayed or non-delayed auditory feedback depending on group assignment) and the Auditory-Motor Adaptation task (production of target words while exposed to formant-shifted auditory feedback that was delayed or non-delayed depending on group assignment). Subjects rated perceived auditory delay following each speech production block. (C) Instrumentation setup for both the Delay Exposure task and the Auditory-Motor Adaptation task (with masking noise used only in the Adaptation task). (D) Time course of the formant-shift feedback perturbation during the Auditory-Motor Adaptation task.

### Delay Exposure task

In the *Delay Exposure* task, subjects produced consonant-vowel-consonant (CVC) words for 30 minutes while hearing their auditory feedback with either a minimal 10 ms delay intrinsic to the involved equipment (labeled “no delay” in the current study; used for the *Control Group* and the two *No-Habitation Groups*), a total delay of 75 ms (10 ms equipment delay plus 65 ms added delay; used for the *Habituation Short-Delay Group*), or a total delay of 115 ms (10 ms equipment delay plus 105 ms added delay; used for the *Habituation Long-Delay Group*). The task was divided into six 5-minute blocks, with blocks alternating between word reading and picture naming (60 trials per block; Figure 1B) to reduce monotony during this relatively long task. For each block, orthographically presented words were drawn randomly from a set of 180 different items or pictorially presented words were drawn randomly from a set of 72 different images.

Subjects’ acoustic speech output was captured with a microphone (SM 58, Shure) positioned 15 cm from the mouth, amplified (DPS II, Applied Research and Technology), routed through a digital vocal processor capable of implementing delays (VoiceOne, TC Helicon, controlled through Musical Instrument Digital Interface [MIDI] signals from a computer workstation), and played back to the subject through insert earphones (ER-3A, Etymotic Research). Immediately prior to each recording session, the intensity of this speech feedback system was calibrated using a 2 cc coupler (Type 4946, Bruel & Kjaer) connected to a sound level meter (Type 2250A Hand Held Analyzer with Type 4947 ½” Pressure Field Microphone, Bruel & Kjaer). During this calibration process, amplification levels were adjusted such that a speaking intensity of 75 dB SPL at the microphone also resulted in an acoustic feedback signal of 75 dB SPL in the insert earphones.

After each block of 60 trials, subjects reported their subjective perception of any delay in the auditory feedback by marking, with a computer mouse, a location along a visual-analog scale presented on a laptop computer monitor (Figure 1B). The scale was presented by means of the Adaptive Visual Analog Scales (AVAS) software program (Marsh-Richard et al., 2009). It was presented as a horizontal 75 mm line across the middle of the screen, with the left and right anchors labeled “No delay” and “Longest delay,” respectively. Mouse clicks along the visual line were automatically recorded by the AVAS software as a numerical score ranging from 0 (extreme left end of the scale) to 100 (extreme right end of the scale). Prior to initiating the *Delay Exposure* task, participants were thoroughly familiarized with this scale by completing a training procedure using the AVAS software. At that time, they were also familiarized with the “No delay” and “Longest delay” anchors of the scale and a point exactly in the middle of the scale. This was accomplished by letting subjects produce 10 CVC words in each of the following three auditory feedback conditions: only the 10 ms hardware-intrinsic delay, a total delay of 150 ms, and a total delay of 75 ms. Thus, the middle of the scale corresponded to the delay experienced by subjects in the *Habituation Short-Delay* (75 ms) group, and the scale extended well beyond the delay experienced by subjects in the *Habituation Long-Delay* (115 ms) group in order to minimize ceiling effects.

Instructions to all subjects during the familiarization phase were as follows: No delay – “What you will hear now is the best that our equipment can do, so you will hear your own speech with no delay,” and after completion of 10 productions “If you hear yourself this way, with no delay in the signal, you should click all the way to the left on the scale where it says no delay”; 150 ms delay – “What you will hear now is the longest delay that could ever happen in our equipment,” and after 10 productions “If you hear yourself with a delay as long as what you just heard, you should click all the way to the right on the scale where it says maximum delay”; 75 ms delay – “What you will hear now will be a delay that is right in the middle between the best and worst possible situations that you just heard,” and after completion of the productions “Could you tell that the delay was in the middle between the best and the worst possible? If you think that the delay that you hear is halfway between the best and worst possible you should click in the middle of the scale”). In this familiarization phase, the middle delay of 75 ms was always presented last, but the order of no delay and the longest delay of 150 ms was varied across subjects.

### Auditory-Motor Adaptation task

The *Auditory-Motor Adaptation* task (Figure 1C) assessed subjects’ speech adjustments to a frequency perturbation in the auditory feedback when this feedback signal was heard with a delay that—depending on group assignment—subjects had or had not been exposed to during the *Delay Exposure* task. The task involved producing 65 epochs of the three monosyllabic words “tech,” “tuck,” and “talk” for a total of 195 productions. The three words were randomized within each epoch and individually presented in the top half of a computer screen in front of the subject. Subjects’ speech was transduced, routed, and played back in the same manner as described above for the *Delay Exposure* task, but pink masking noise was also mixed into the earphones at 68 dB SPL to minimize the availability of non-manipulated bone-conducted feedback prior to onset of the delayed earphones signal (see Max & Maffett 2015 for more details on the required masking level). Subjects were aided in maintaining a consistent speaking level by presenting color-coded visual feedback about speech intensity in the bottom half of the computer screen, with a target level between 72 and 78 dB SPL as measured at the microphone (15 cm from the mouth).

The frequency perturbation in the adaptation task consisted of an increase in the frequency of all formants (i.e., vowel resonances) to a maximum shift of +2.5 semitones, a manipulation that has been shown consistently to induce speech auditory-motor adaptation if implemented in real-time without delay (Daliri & Max, 2018; Max & Maffett, 2015; Max, Wallace, & Vincent, 2003). The formant shift implementation followed the same time course for all participants (Figure 1D): 30 trials with unaltered feedback (*baseline* phase); 60 trials during which the formant frequencies were incrementally increased to a maximum shift of +2.5 semitones (*ramp shift* phase); 60 trials during which the feedback shift was maintained at +2.5 semitones (*full shift* phase); and 45 trials after unaltered feedback had been restored (*after effects* phase or washout phase). Depending on specific group assignment, this formant-shifted auditory feedback signal was presented into the subject’s earphones with one of the delays used during the *Delay Exposure* task: no delay (other than the 10 ms equipment delay), a total delay of 75 ms, or a total delay of 115 ms. After completion of the *Auditory-Motor Adaptation* task, all subjects provided one final rating of their subjective perception of feedback delay using the visual analog scale described above.

### Data Processing and Extraction

Subjects’ speech was digitized directly onto a computer (16-bit, 44.1 kHz) and then analyzed offline using custom software that combines routines from Praat (Version 6.0.39; Boersma et al., 2018) and Matlab (The Mathworks, Natick, MA). Specifically, we extracted the first and second formant frequencies (F1 and F2), averaged over the middle 20% of each trial’s vowel portion, by means of Praat’s default linear predictive coding algorithm (Boersma et al., 2018). In rare cases of speaking errors or formant tracking difficulties, the F1 and F2 values for missing trials were replaced by estimates linearly interpolated from neighboring productions of the same word. The first three baseline trials for each word were discarded to prevent contamination of the subsequent processing step (frequency normalization) by relatively large F1 changes that often occur when participants adjust their articulatory movements to bring output intensity into the desired range indicated by visual feedback (a phenomenon of particular concern when initial trials are produced more loudly due to the presence of masking noise in the earphones).

The extracted F1 and F2 frequencies in Hertz (Hz) were then normalized within each speaker by transforming each trial to *cents* units (100 cents = 1 semitone) using the following formula:

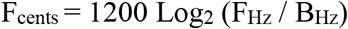

where F_Hz_ corresponds to the trial’s formant frequency in Hz, and B_Hz_ corresponds to subject’s baseline formant frequency in Hz for the same word (calculated as the median of the subject’s 7 analyzed productions of that word in the *baseline* phase). Lastly, each subject’s overall change in acoustic speech output in response to the auditory feedback manipulation was quantified by computing an *adaptation index* from the normalized formant frequencies for the last 15 trials (5 trials of each word) in the *full shift* phase of the adaptation task. This adaptation index was based on data averaged across F1 and F2 and across the three test words.

### Statistical Analyses

#### Perceptual ratings of feedback delay

To compensate for any potential changes in delay perception that occur even in the absence of feedback delay (i.e., drift over time), we first normalized the perceptual ratings from each participant in the four experimental groups by subtracting, at each of the seven time points, the corresponding average responses of the no-delay *Control Group*. Thus, the final perceptual data set consisted of four groups’ normalized ratings after each 5-minute block of speech production during the *Delay Exposure* task (ratings #1 through #6) and immediately after the *Auditory-Motor Adaptation* task (rating #7). Importantly, the questions under investigation cannot be addressed by examining overall group or rating time point differences. Rather, five specific *a priori* pair-wise comparisons are needed to determine (a) whether participants who already experienced feedback delay during the exposure task did indeed show a decrease in their perception of the extent of delay when completing the auditory-motor adaptation task (rating #7 vs. #1 for the combined habituation groups); (b) whether these participants in the habituation groups completed the adaptation task while remaining at the same level of habituation as achieved at the end of the preceding exposure task (ratings #7 vs. #6 for the combined habituation groups); (c) whether participants who did not experience feedback delay during the exposure task did in fact increase their ratings when such delay was experienced in the auditory-motor adaptation task as compared with both the beginning and end of the preceding exposure task (ratings #7 vs. #1 and #7 vs. #6 for the combined no-habituation groups); and (d) whether the participants who experienced feedback delay only in the adaptation task perceived it to be of similar duration as reported at the beginning of the exposure task by those who were being pre-exposed at that time (rating #7 for the combined no-habituation groups vs. rating #1 for the combined habituation groups). In the below Results section, all listed *p* values are the adjusted values obtained by applying the Holm-Bonferroni correction method (Holm, 1979) for interpretation against an overall α level of .05 for this family of five tests.

#### Auditory-motor adaptation

Adaptation to the formant-shift perturbation manifests itself as a change in the produced vowel formants relative to the baseline phase during which only unperturbed feedback was heard. Therefore, the analysis approach required first asking whether or not adaptation did actually occur when feedback was presented with no delay, with a 75 ms delay, and with a 115 ms delay (i.e., whether or not changes from baseline were statistically significant separately within the *Control Group*, the *No-Habituation Short-Delay Group*, and the *No-Habituation Long-Delay Group*). Next, it was necessary to conduct two between-group *a priori* pair-wise comparisons to test the effect of feedback delay in isolation by asking whether the amount of adaptation in either the *No-Habituation Short-Delay Group* or the *No-Habituation Long-Delay Group* was different from the amount observed for the *Control Group* (these three groups differed in delay experienced during the adaptation task but had not experienced delay in the exposure task). Lastly, the effect of prior habituation could be tested in isolation with two more between-group *a priori* pair-wise comparisons that asked whether the amount of adaptation in the *Habituation Short-Delay Group* and *Habituation Long-Delay Group* was different from that observed in the *No-Habituation Short-Delay Group* and the *No-Habituation Long-Delay Group*, respectively. We considered these statistical tests on the adaptation data as one family of three within-group comparisons and one group of four between-group comparisons, and all *p* values listed below are Holm-Bonferroni adjusted values that are interpreted against a family-wise error rate of α = .05.

## Results

### Perceptual ratings of feedback delay

The four experimental group’s normalized ratings after each 5-minute block of speech production during the *Delay Exposure* task (judgments #1-6) and immediately after the *Auditory-Motor Adaptation* task (judgment #7) are shown in Figure 2. The two groups that did experience a feedback delay during the *Delay Exposure* task (i.e., the two habituation groups) showed a gradual decrease in their judgment of the extent of the delay. In fact, although there was large inter-subject variability (see also Yamamoto & Kawabata, 2011), the average rating after just four blocks of speaking indicated no perceived delay at all (i.e., a rating of 0 ms relative to the *Control Group* that rated the same feedback signal without exposure to a delay). Thus, subjects in these two groups showed clear evidence of perceptual habituation to the delay across the repeated blocks of speaking, at least in terms of the group average rating. Importantly, the final ratings of both groups indicated that they completed the *Auditory-Motor Adaptation* task also in a perceptually habituated state. The reduction in perceived delay between the first and seventh ratings was statistically significant [*t*(21) = 3.483, *p* = .009] whereas the difference between ratings six and seven was not [*t*(21) = 0.660, *p* = .516].

**Figure 2.**
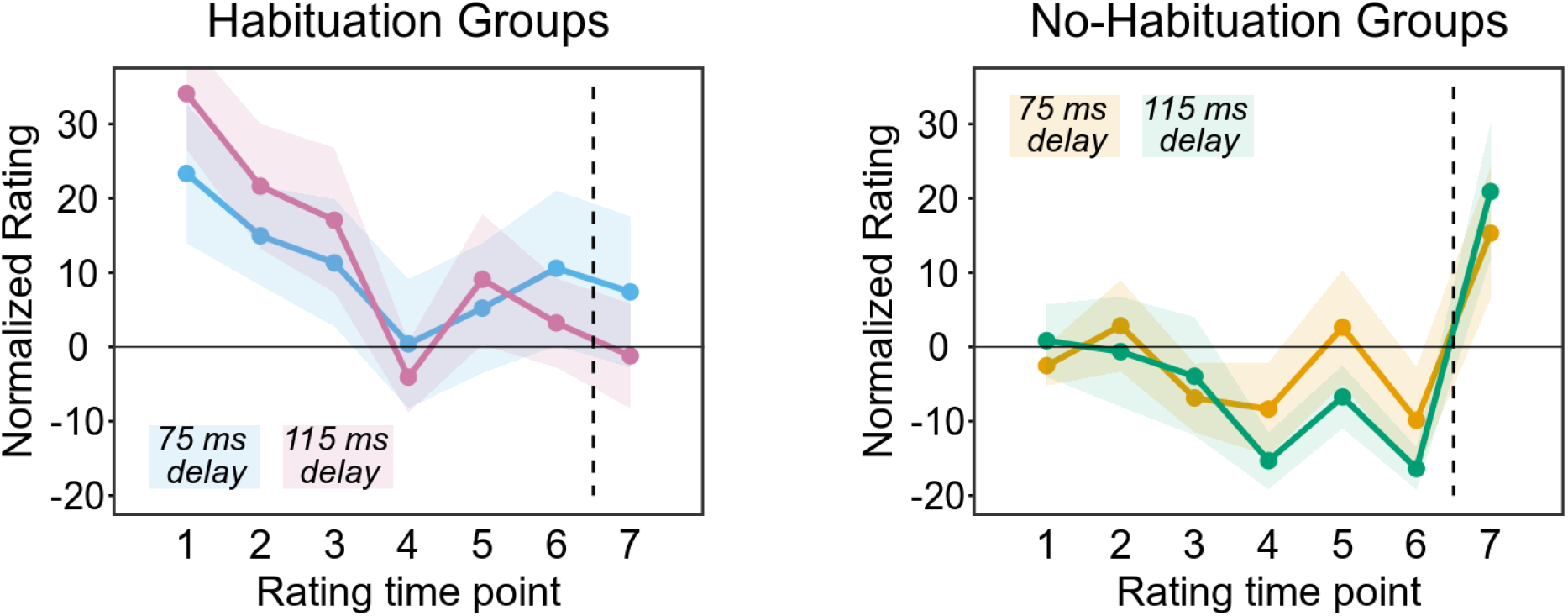
Group means (normalized by subtracting the average responses of the no-delay Control Group) for ratings of perceived auditory feedback delay by the subject groups who experienced a delay throughout the Delay Exposure task (habituation groups, left panel), and the subject groups who experienced no delay during the Delay Exposure task (no-habituation groups right panel). Ratings at time points 1 through 6 were completed during the Delay Exposure task (each rating following a 5-minute block of speaking) whereas the rating at time point 7 was completed immediately following the Auditory-Motor Adaptation task. Shaded areas show +/− one standard error of the mean.

On the other hand, the subject groups that experienced no delay during the *Delay Exposure* task (the two no-habituation groups) showed little change in their ratings across the initial six speaking blocks and then an increase in their judgment of the delay for the adaptation task (which was the first time these groups were exposed to a delay). Thus, the latter subjects completed the adaptation task in a non-habituated state. Their perceived delay at the seventh rating point was statistically significantly longer than that at both the first rating [*t*(21) = −3.496, *p* = .009] and the sixth rating [*t*(21) = −4.791, *p* < .001]. Moreover, these no-habituation subjects’ perceived delay at the seventh rating point (addressing their perception during the adaptation task) was not statistically significantly different from that of the habituation subjects at the very first rating point (when initially exposed to the delay) [*t*(42) = 1.227, *p* = .453].

### Auditory-Motor Adaptation task

Data for the no-delay *Control Group,* the *No-Habituation Short-Delay Group*, and the *No-Habituation Long-Delay Groups* are shown in Figure 3. The *Control Group,* which experienced no delay during either the *Delay Exposure* or *Auditory-Motor Adaptation* tasks, started decreasing their produced formants during the ramp shift phase, further decreased these formants throughout the full shift phase, achieved a maximum amount of adaptation of 114 cents (corresponding to 46% of the magnitude of the implemented perturbation), and then gradually returned toward baseline during the after-effects phase, but did not completely reach baseline performance before the end of the task. This pattern of adaptive changes is fully consistent with previous studies using the same auditory perturbation (Max & Maffett, 2015; Max, Wallace, & Vincent, 2003). Using a one-sample *t*-test comparing the averaged F1 and F2 frequencies at the end of the full shift phase to zero (i.e., the average baseline value), the *Control Group*’s amount of adaptation was statistically significant [*t*(10) = −8.727, *p* < .001]. Overall, the *No-Habituation Short-Delay Group* and the *No-Habituation Long-Delay Group* showed formant production changes in the same direction, and the final amount of adaptation at the end of the full shift phase was also statistically significant for both groups [75 ms group: *t*(10) = −4.023, *p* = .004; 115 ms group: *t*(10) = −4.112, *p* = .004].

**Figure 3.**
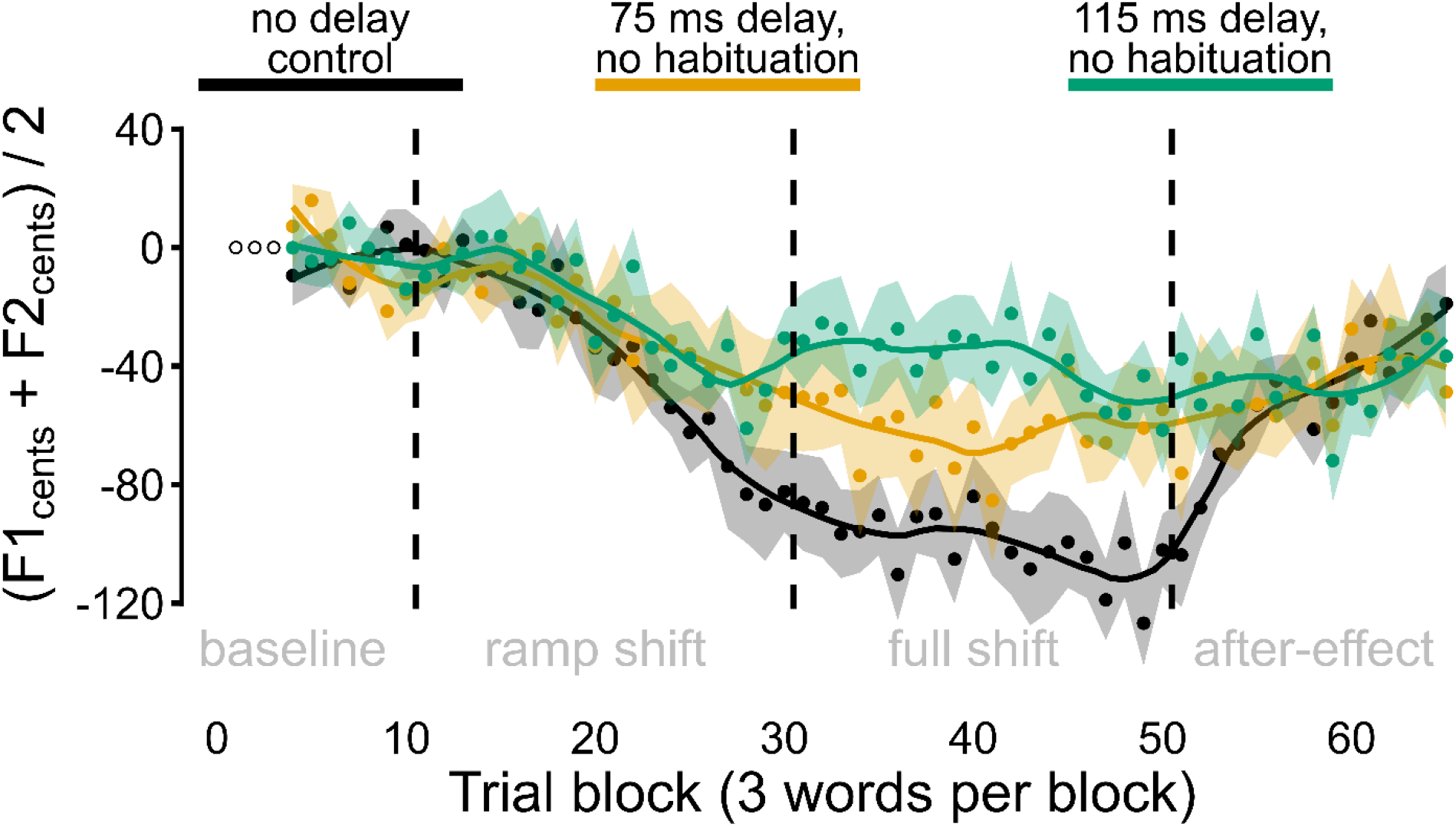
Speech auditory-motor adaptation to a ramped perturbation that consisted of an upward shift of vowel formant frequencies in the auditory feedback signal (250 cents in the full shift phase). Adaptive changes in the averaged Formant 1 (F1) and Formant 2 (F2) of subjects’ productions are calculated in cents relative to the baseline phase during which no formant shift was applied. Data are shown for the Control Group that experienced no feedback delay during either 30 minutes of speaking or the subsequent adaptation task, the No-Habituation Short-Delay Group that experienced no feedback delay during 30 minutes of speaking but a 75 ms feedback delay during the subsequent adaptation task, and the No-Habituation Long-Delay Group that experienced no feedback delay during 30 minutes of speaking but a 115 ms feedback delay during the subsequent adaptation task. Data points represent the group mean formant frequency values for each block of the 3 different test words (averaged across F1 and F2); the first 3 blocks of trials were discarded; shaded regions show standard errors of the mean; solid lines are loess smoothed fits (span .25).

However, as is clear from Figure 3, the amount of adaptation was negatively impacted when a delay was added to the auditory feedback signal. Whereas the final amount of adaptation in the *Control Group* reached 114 cents, in the *No-Habituation Short-Delay* and *No-Habituation Long-Delay* groups it reached only 59 cents and 49 cents, respectively. Using Welch *t*-tests (with adjusted degrees of freedom to account for unequal group variances and with the adjusted *p* values from the Holm-Bonferroni correction), this reduction in auditory-motor adaptation relative to the *Control Group* was statistically significant for both the *Short-Delay Non-Habituation Group* [*t*(19.75) = −2.816, *p* = .032] and the *No-Habituation Long-Delay Group* [*t*(19.84) = −3.675, *p* = .006].

Lastly, the effect of 30 minutes of prior feedback delay habituation on the amount of auditory-motor adaptation to the formant shift perturbation was examined for the 75 ms and 115 ms time delays. Figure 4 shows the adaptation time course for the *No-Habituation Short-Delay Group* vs. the *Habituation Short-Delay Group* (top panel) and for the *No-Habituation Long-Delay Group* vs. the *Habituation Long-Delay Group* (bottom panel). Given that one might predict prior delay habituation to reverse the negative effect of such delays on auditory-motor adaptation, the time course of produced formant changes in the no-delay *Control Group* is also shown again for comparison. Despite the fact that 30 minutes of delay exposure had significantly reduced or eliminated the subjective perception of delay in the habituation groups (see perceptual rating results above), these subjects showed no improvement at all in their amount of auditory-motor adaptation as compared with the subjects who had not been previously exposed to the delay. Welch *t*-tests (again with adjusted degrees of freedom and with Holm-Bonferroni adjusted *p* values) confirmed the absence of such an effect for both the 75 ms condition [*t*(19.71) = −0.063, *p* = 1.00] and the 115 ms condition [*t*(19.26) = −0.345, *p* = 1.00]. Thus, prior habituation to the auditory feedback delay did not reduce the negative impact of this delay on the magnitude of the adaptation effect.

**Figure 4.**
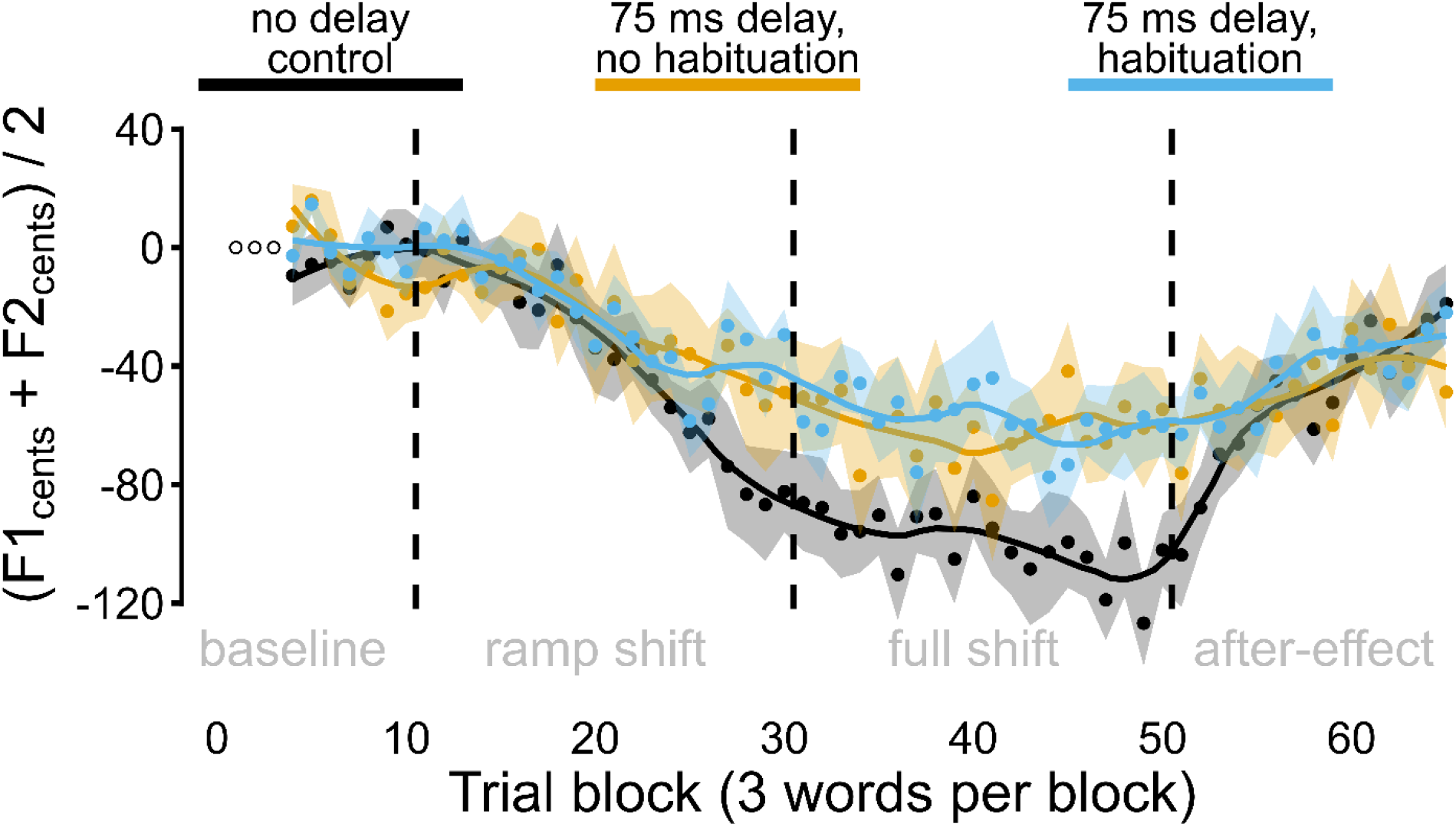
Speech auditory-motor adaptation to a ramped upward shift (250 cents) of vowel formant frequencies in the auditory feedback signal. A: Comparison of the Control Group that experienced no feedback delay during either 30 minutes of speaking or the subsequent adaptation task, the No-Habituation Short-Delay Group that experienced no auditory feedback delay during 30 minutes of speaking but 75 ms delay during the subsequent adaptation task, and the Habituation Short-Delay Group that experienced 75 ms auditory feedback delay during both 30 minutes of speaking and the subsequent adaptation task. B: Comparison of the same Control Group with the No-Habituation Long-Delay group (no auditory feedback delay during 30 minutes of speaking, 115 ms delay during the adaptation task) and the Habituation Long-Delay Group (115 ms delay during both 30 minutes of speaking and the subsequent adaptation task). Data points indicate group mean formant frequencies for each block of the 3 different test words (averaged across F1 and F2); the first 3 blocks of trials were discarded; shaded areas indicate standard errors of the mean; solid lines are smoothed fit loess functions with span .25.

**Figure 5.**
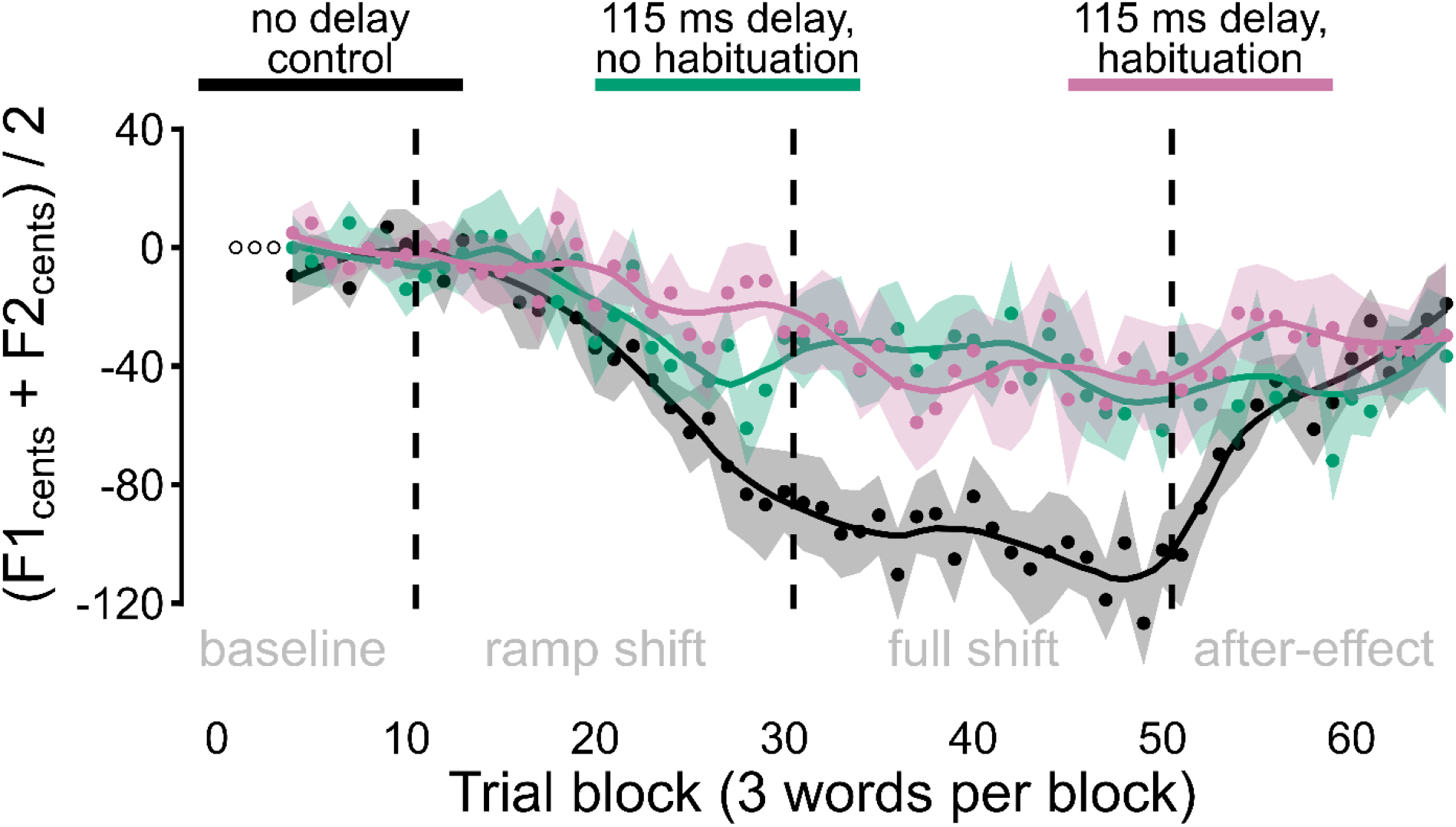

## Discussion

Prior studies on reaching movements have shown that prism or visuomotor rotation tasks completed with visual feedback delays of 100-5000 ms yield reduced but robust motor learning effects (i.e., changes in reach direction during the perturbation phase followed by persisting after-effects during the wash-out phase) (Honda et al. 2012a; Kitazawa et al. 1995; Tanaka et al. 2011). Moreover, related studies have shown that the limited negative effect of a 200 ms visual delay on visuomotor adaptation can be further reduced through prior exposure and habituation to the delay (Honda 2012a, 2012b). For speech movements, on the other hand, prior studies have suggested that even delays as short as 100 ms nearly or completely eliminate auditory-motor adaptation to formant-shifted feedback (Max & Maffett 2015; Mitsuya et al. 2017), and the potential effects of prior delay habituation remained entirely unknown. We therefore investigated whether perceptual habituation to an auditory feedback delay would mitigate the negative effects of such delays in a speech auditory-motor adaptation task.

In an initial speech production task without formant shift perturbation, two groups of participants spoke for 30 minutes with either short or long feedback delays (75 ms or 115 ms total delay for the *Habituation Short-Delay Group* and the *Habituation Long-Delay Group*, respectively). Participants in both groups showed strong evidence of habituating to the delay, judging the feedback delay to get gradually shorter throughout this 30-minute *Delay Exposure* task and to have almost entirely disappeared by the end of the speaking block. In the subsequent *Auditory-Motor Adaptation* task, the subjects from these two groups remained perceptually habituated to the delay while a formant-shift perturbation was also incrementally added to the auditory feedback signal. Interestingly, these two delay-habituated groups showed no benefit at all in adapting to the formant-shift perturbation as compared with two other, non-habituated groups who had not been previously exposed to the 75 or 115 ms feedback delays that they experienced during the adaptation task. Thus, although we found that perceptual habituation does occur for feedback delays in speech production, we found no evidence for a facilitating effect of such habituation on speech auditory-motor adaptation to a formant-shift perturbation in the presence of feedback delays. All four subject groups that completed the formant-shift adaptation task with a feedback delay (two habituated groups and two non-habituated groups) showed only 40-50% of the extent of adaptation seen in the *Control Group* that completed both the initial 30-minute speaking block and the subsequent adaptation task without any delay added to the auditory feedback. Thus, findings related to the overall extent of adaptation do again confirm that even relatively short feedback delays have substantial detrimental effects on speech auditory-motor adaptation.

It is worth noting that, although delays did have this negative effect on adaptation, the final extent of adaptation achieved by the two 75 ms delay groups as well as the two 115 ms delay groups was larger than that observed in two previous formant perturbation studies with 100 ms delay (Max & Maffett 2015; Mitsuya et al. 2017). Thus, the detrimental effect of auditory feedback delay on speech auditory-motor adaptation was less severe here than in previous studies. The reasons for this finding are unclear, but one major methodological difference is that only the participants from the present study first completed a separate block of 30 minutes of speaking before starting the adaptation task, with the exact same instrumentation used for routing auditory feedback in both tasks (recall that the initial speaking task for all participants served to create separate groups that had or had not habituated to the feedback delay, but no formant perturbation was present for any of the groups). Consequently, the extra half hour of speaking in this set-up may have helped participants with getting used to the different sound quality associated with hearing their own speech through insert earphones as opposed to free field air conduction and bone conduction as in typical daily-life speaking conditions. The prior acoustic familiarization, in turn, may have resulted in the neural control system being more sensitive to additional auditory error caused by the subsequent formant perturbation. This interpretation is supported by the present study’s finding that the *Control Group* also showed a final extent of adaptation that is larger (close to 50% of the implemented formant perturbation) than that typically observed in speech production experiments with formant shift perturbations.

Also noteworthy is the partial overlap of the learning curves for the *No-Habituation Short-Delay Group* (no prior delay habituation, 75 ms delay adaptation task) and the *No-Habituation Long-Delay Group* (no prior delay habituation, 115 ms delay adaptation task). In fact, these two delay groups showed very similar learning trajectories during the ramp portion of the perturbation phase before the curves diverged for most of the hold portion of the perturbation phase. It could therefore be speculated that a longer delay of 115 ms is tolerated just as well as a shorter delay of 75 ms as long as other sensory prediction errors (here related to experimental manipulation of the heard formant frequencies) are small, but that the combination of longer delay *plus* increasingly large prediction errors results in a disruption of the ongoing learning. Under such circumstances, credit assignment for the growing overall error may be shifted to external sources rather than self-generated actions. However, this speculative idea is hard to reconcile with the observation that the corresponding habituated participants (*Habituation Long-Delay Group*) did not show a similar initial learning trend during the ramp portion of the perturbation phase.

Crucially, our central finding that prior habituation to auditory delays of 75 or 115 ms does not mitigate the detrimental effects of auditory feedback delay on speech auditory-motor learning stands in contrast with Honda’s (2012a, 2012b) reports that habituation to visual delays of 260 ms significantly reduces the negative effects of visual feedback delay on visuomotor learning for reaching movements. Our result is more in line with the earlier conclusion by Tanaka et al. (2011), based on prism adaptation for pointing movements with 136 ms visual delay, that “Physical delay but not subjective delay determines learning rate […]” (p. 257). This is an intriguing result given that our methodology was much more similar to that used by Honda (2012a, 2012b) rather than that of Tanaka et al. (2011): (a) our delay exposure task involved 360 trials (100-120 trials in Honda [2012a, 2012b] vs. only 60 trials in Tanaka et al. [2011]), (b) our auditory formant perturbation was introduced incrementally over many trials (visual perturbations were introduced incrementally in Honda [2012a, 2012b] vs. abruptly in Tanaka et al. [2011]), and (c) we presented auditory feedback throughout the entire word production for each trial (visual feedback reflected the full motion path in Honda [2012a, 2012b] vs. only movement endpoint in Tanaka et al. [2011]).

Hence, the effect of prior delay habituation on sensorimotor learning may be both effector system specific and situation specific. Here, we found that delay habituation had no benefit at all for subsequent speech auditory-motor adaptation with auditory feedback delays even in conditions that had been found to yield a clear delay habituation benefit in the case of reaching movement visuomotor adaptation (that is, conditions in which full feedback was available during each trial and the perturbation was introduced incrementally across trials). Such differences between speech and limb motor learning in feedback delay sensitivity and delay perception plasticity also warrant further research on the generalizability of theoretical perspectives regarding delay representation in neural systems (e.g., the state representation model recently proposed by Avraham et al. 2017) across different sensorimotor domains. Within-subjects analyses of speech and limb motor learning tasks may prove particularly valuable in this regard.

For example, it is plausible that, in speech motor control, heightened sensitivity to sensory feedback delays and increased resistance to delay habituation result from fundamental differences in this system’s temporal constraints on sensorimotor integration and movement planning. Speech articulation requires the coordination of numerous biomechanical structures distributed across and within different effectors (lips, tongue, jaw, velum, larynx) and achieves rates up to 5-6 syllables (or 10-15 individual sounds) per second (Fonagy and Magdics 1960; Levelt 1989; Zemlin 1998). Speech movement durations are often as short as 50-200 milliseconds, and movement amplitudes are as small as a few millimeters (Gracco 1994; Max et al. 2003; Ostry and Munhall 1985). Additionally, as a second specific characteristic, the ultimate goals of articulatory movements are sequences of sounds that are intelligible to a listener, and the movements are planned, at least in part, in terms of those acoustic targets (Callan et al. 2000; Feng et al. 2011; Guenther 1994; Guenther et al. 1998, 1999; Lindau et al. 1972; Perkell et al. 1997, 2000). The latter point emphasizes the unique need for the neural controller for speech to take into account not only dynamic and kinematic transformations similar to those involved in limb movements but also additional complex transformations from vocal tract tube shapes and constrictions to the acoustic speech output (Fant 1980; Stevens 2000). Despite subjects’ demonstrated capacity for perceptual delay habituation of the acoustic output signal, limitations related to the *physical* delay prevent the neural controller for speech from adjusting sensory predictions to take account of the effect of delay across all the input-output transformations or to increase its reliance on the delayed feedback for adjusting motor commands. Given that habitation did occur, we speculate that the involved sensory predictions were, in fact, appropriately adjusted during the initial delay exposure phase, but that this process affecting participants’ *awareness* of the delay nevertheless failed to drive motor learning during the formant perturbation phase.

Considering that our study specifically manipulated participants’ awareness of the delay (leading to no awareness of delay in the habituated groups), the lack of an effect of delay habituation on speech auditory-motor learning suggests another potential factor that may contribute to the different results obtained here for speech auditory-motor learning as opposed to prior work on reach visuomotor learning (Honda 2012a, 2012b). Whereas reach visuomotor adaptation involves a combination of both implicit learning and explicit strategy use (Anguera et al. 2010; Fernandez-Ruiz et al. 2011; Holland et al. 2018; McDougle et al. 2016; Taylor et al. 2014), multiple lines of evidence suggest that speech auditory-motor adaptation to formant perturbations likely constitutes an almost entirely implicit form of learning. First, naïve participants have no explicit knowledge of the complex relationship between vocal tract configurations and vowel formant frequencies; thus, any explicit strategy use itself would first need to be acquired through extensive trial-and-error learning. Second, studies on speech sensorimotor learning have demonstrated that subjects are unaware that they make any changes in their speech output in the presence of the perturbation (Kim and Max 2020), and that there is no difference in the extent of adaptation when subjects are instructed to ignore the feedback or to avoid compensating for the perturbation (Keough et al. 2013; Munhall et al. 2003). Third, the explicit component of limb visuomotor adaptation is negatively affected by simultaneous speech auditory-motor adaptation, but there is no interference of the limb learning task on the speech learning task (Lametti et al. 2020). Combining those findings with recent work indicating that delayed feedback negatively affects implicit learning without much impact on explicit strategy selection (Brudner et al. 2016; Schween and Hegele 2017), we suggest that both the strong negative impact of feedback delay on speech auditory-motor learning and the lack of a beneficial effect of prior delay habituation may be due to this specific form of sensorimotor learning depending mostly or exclusively on implicit learning processes. In particular, we hypothesize that the inherently implicit nature of speech auditory-motor learning prevents delay-habituated participants from compensating through explicit strategy use when they have no awareness of the delay yet continue to experience large discrepancies between the intended and achieved auditory targets (as the physical delay continues to impair their implicit learning). By purposefully designing new speech auditory-motor learning paradigms that depend to varying degrees on implicit versus explicit learning, future studies might be able to directly test this hypothesis.

In sum, the present study identified a substantial decrease in speech auditory-motor learning when auditory feedback was delayed by 75 or 115 ms. Even though a preceding half hour of delay exposure was found to be sufficiently long to induce perceptual habituation to the delay, such prior exposure yielded no benefit in a subsequent adaptation task in which a formant shift perturbation was added to the delayed auditory feedback signal. Thus, we conclude from the clear dissociation of the perceptual data and the sensorimotor adaptation data that such short-term habituation to auditory feedback delay is not effective in reducing the negative impact of delay on speech auditory-motor adaptation. We further hypothesize that this finding likely results from fundamental differences in control requirements for the speech and reach effector systems or from different contributions of implicit and explicit learning mechanisms when these effector systems adapt to perturbed feedback signals.

## Acknowledgments

This research was supported by grants R01DC014510 and R01DC017444 from the National Institute on Deafness and Other Communication Disorders, grant MOP-137001 from the Canadian Institutes of Health Research, and the Natural Sciences and Engineering Research Council (NSERC-Canada). The content is solely the responsibility of the authors and does not necessarily represent the official views of the National Institute on Deafness and Other Communication Disorders, the National Institutes of Health, or the Canadian Institutes of Health Research.

## Footnotes

1 It has been shown that visual feedback delays of 75 or 150 ms during reaching movements had no statistically significant effect on adaptation to a force field (i.e., a somatosensory perturbation), but in that case even the complete absence of visual feedback did not significantly disrupt learning (McKenna et al., 2017).

